# Technical Report: Targeted Proteomic Analysis Reveals Enrichment of Atypical Ubiquitin Chains in Contractile Murine Tissues

**DOI:** 10.1101/2020.03.18.983312

**Authors:** Tiaan Heunis, Frederic Lamoliatte, José Luis Marín-Rubio, Abeer Dannora, Matthias Trost

**Author notes:** These authors contributed equally to this work (Tiaan Heunis, Frederic Lamoliatte). Corresponding author: Matthias Trost.

## Abstract

Ubiquitylation is an elaborate post-translational modification involved in all biological processes. Its pleotropic effect is driven by the ability to form complex polyubiquitin chain architectures that can influence biological functions. In this study, we optimised sample preparation and chromatographic separation of Ubiquitin peptides for Absolute Quantification by Parallel Reaction Monitoring (Ub-AQUA-PRM). Using this refined Ub-AQUA-PRM assay, we were able to quantify all ubiquitin chain types in 10-minute LC-MS/MS runs. We used this method to determine the ubiquitin chain-linkage composition in murine bone marrow-derived macrophages and different mouse tissues. We could show tissue-specific differences in ubiquitin levels in murine tissues, with polyubiquitin chain types contributing a small proportion to the total pool of ubiquitin. Interestingly, we observed enrichment of atypical (K33) ubiquitin chains in heart and muscle. Our approach enabled high-throughput screening of ubiquitin chain-linkage composition in different murine tissues and highlighted a possible role for atypical ubiquitylation in contractile tissues.

## Main Text

Ubiquitylation is a post-translational modification that results from the addition of ubiquitin, a 76 amino acid protein, to generally the ε-amino group of a lysine residue. Ubiquitylation plays a role in practically all eukaryotic biological processes. It is therefore not surprising that aberrant ubiquitylation is linked to several human diseases, including cancer, infectious, neurodegenerative, and developmental diseases [1]. Protein substrates can be modified by monoubiquitin or can be polyubiquitylated, which is formed by further ubiquitylation of any lysine (K6, K11, K27, K29, K33, K48, and K63) or the N-terminal methionine (M1) of ubiquitin. This allows formation of complex chain architectures (homotypic, heterotypic, or branched chains), which can ultimately influence a substrates fate [2]. However, the biological functions of atypical and mixed ubiquitin chains are only beginning to emerge [1,3]. Therefore, methods are needed to rapidly screen the ubiquitin chain-linkage landscape in cells and tissues to advance our understanding of ubiquitin signalling, especially for atypical linkage types as there are few tools available that facilitate their analysis [4]. Recent technological advances in mass spectrometry-based proteomics have enabled quantification of endogenous levels of ubiquitin chain types by selected reaction monitoring (Ub-AQUA-SRM) [5]. This approach has subsequently been adapted to quadrupole-Orbitrap instruments using parallel reaction monitoring (PRM) [6] to determine chain-linkage composition of purified substrates [7], whole-cell lysates [8], to quantify K48-K63 branched chains [9,10], to gain information on higher-order chain-linkage architecture of ubiquitylated substrates [7] and to quantify phosphorylated and acetylated ubiquitin [11,12]. Here we further refined the Ub-AQUA-PRM method for absolute quantification of ubiquitin chain types. We optimised peptide oxidation, ion-pairing agents, and implemented microflow chromatographic separation for increased assay sensitivity and throughput. Our refined Ub-AQUA-PRM assay is ideal for high-throughput screening of ubiquitin chain types in diverse samples, as demonstrated here using murine primary cells and tissues.

Methionine-containing peptides can be challenging to analyse by PRM-based methods, as they can occur in different oxidation states [13]. This can become problematic for quantification when the peptide signal is divided across multiple oxidation states or, importantly, when there are differences between the endogenous peptide and the spike-in standard. Current Ub-AQUA-PRM strategies analyse the methionine sulfoxide version of methionine-containing peptides, induced by using low concentrations of H_2_O_2_ at 4°C [13,14]. However, these peptides are stable for only 21.5 hours, before oxidation state shifts are observed [13]. We therefore fully oxidised methionine residues into the sulfone derivative using H_2_O_2_. No difference in methionine sulfone-containing peptide (M1 or K6) intensity was observed without H_2_O_2_ at room temperature or 60°C for 2 hours (Supplementary Figure 1a, b). However, a methionine sulfone was generated when using 0.1% H_2_O_2_ at 60°C for 2 hours, but the sulfoxide version still contributed substantially to M1 and K6 peptide intensity. Both methionine-containing peptides (M1 and K6) were converted into a stable methionine sulfone by more than 99.9%, based on peptide intensity, when using 1% H_2_O_2_ for 2 hours at 60°C (Supplementary Figure S1a, b). The other peptides analysed were unaffected under the same conditions (Supplementary Figure S1c). This indicates that our oxidation procedure does not introduce undesirable artifacts. We, therefore, used 1% H_2_O_2_ for 2 hours at 60°C for peptide oxidation in all further analyses. Next, we determined the optimal normalised collision energy (NCE) required for fragmenting each peptide. Detailed information on NCE used for each peptide is available in Supplementary Table S1. Additionally, we also tested the effect of two commonly used ion-pairing agents in reversed-phase chromatography for their effect on peptide retention and ion intensity, namely formic acid (FA) and trifluoroacetic acid (TFA). We compared FA and TFA at concentrations ranging from 0.2 to 5.0% (v:v) in the loading buffer (Supplementary Figure S2). For all these injections, buffer A and B remained the same. The K29 peptide is the most hydrophilic of the ubiquitin peptides, but we did not observe a major difference in peak intensity or peak shape between 5.0% FA and 0.2% TFA for this peptide (Supplementary Figure S2a). However, we observed a marked decrease in intensity for other ubiquitin peptides when using TFA, even as low as 0.2% (Supplementary Figure S2b). We, therefore, used 5.0% FA in the sample buffer for all further analyses. Finally, we determined the lower limit of detection (LLOD) and lower limit of quantification (LLOQ) of each peptide in a simple (ubiquitin peptides only) and complex (*E. coli* tryptic digest) matrix. We opted for an *E. coli* tryptic digested as spike-in matrix as no endogenous ubiquitin is present in this organism that would interfere with the assay. LLOD for peptides were as low as 0.5 amol on column. In the simple matrix we could achieve LLOQ ranging from 50 amol for the M1 and K29 peptides to 1.5 fmol for the K11, K63 and TITTITLEVEPSDTIENVK ubiquitin peptides. Similar results were obtained for ubiquitin peptides in the complex matrix (Supplementary Figure S3), reaching LLOQ as low as 0.1 fmol/μg protein injected. These results indicate that our refined Ub-AQUA-PRM assay could accurately quantify low levels of endogenous ubiquitin chain types in complex matrixes.

**Figure 1.**
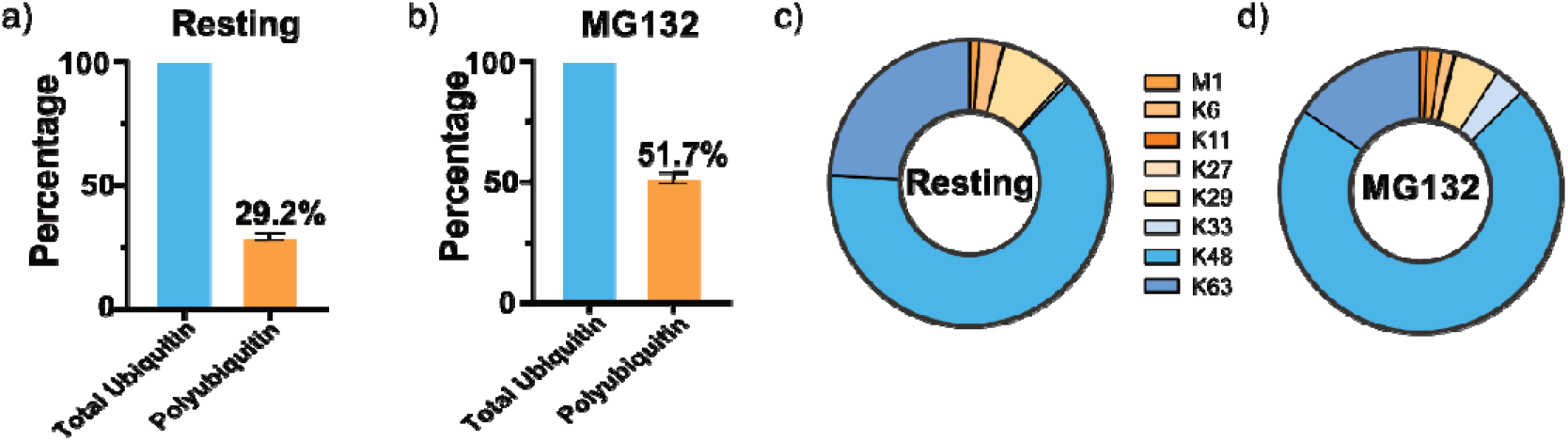
All eight polyubiquitin chain types are present in murine bone marrow-derived macrophages. Percentage of polyubiquitin chain types present in resting, unstimulated (a) and MG132-treated (b) bone marrow-derived macrophages, compared to the total pool of ubiquitin. Contribution of each of the eight different ubiquitin chain types to the total pool of polyubiquitin in resting, unstimulated (c) and MG132-treated bone marrow-derived macrophages (d). Error bars indicate standard deviation.

We used our Ub-AQUA-PRM assay to screen the ubiquitin chain-linkage composition in primary murine macrophages (BMDMs), given the importance of ubiquitin in innate immunity [15–18]. Additionally, we validated our Ub-AQUA-PRM assay using MG132, a proteasome inhibitor that results in ubiquitin accumulation, especially K48 chains [19,20]. An increase in the percentage of polyubiquitin chains was observed in BMDMs after treatment with MG132, as would be expected (Figure 1a, b). All linkage types could be detected in murine BMDMs with K48 (63.2%), K63 (24.2%) and K29 (8.0%) contributing 95.4% of the total ubiquitin chain-linkage composition of whole-cell lysates (Figure 1c). These results are in agreement with previous reports in cell lines [7,21]. Similarly, K48 (71.8%), K63 (15.5%) and K29 (5.2%) contributed 92.5% of the total ubiquitin chain-linkage composition of BMDMs during MG132 treatment (Figure 1c). Next, we determined whether there was any difference in ubiquitin chain types during treatment with MG132 for 6 hours. We observed a statistically significant increase in the majority of ubiquitin chain types, including total ubiquitin, during MG132 treatment (Supplementary Figure S4), which is in agreement with previous reports [7,19,22]. However, K11 and K27 chain types were below the limit of quantification during our analysis. These results indicate that K48, K63 and K29 linkage types dominate the ubiquitin chain-linkage compositions in murine BMDM whole-cell lysates and that up to ~29.2% of total ubiquitin is in the polyubiquitylated form in resting BMDMs.

Lastly, we used our Ub-AQUA-PRM assay to screen the ubiquitin chain-linkage composition of six different murine tissues (brain, heart, kidney, lung, muscle, and spleen) from three different mice. We were able to detect ubiquitin in all samples, with the brain, kidney, and spleen having the highest levels of total ubiquitin (Supplementary Figure S5). A statistically significant difference in total ubiquitin was observed in the brain, which had higher levels of total ubiquitin than the heart, lung, and muscle (Supplementary Figure S5). Interestingly, only a small percentage of total ubiquitin seems to be polyubiquitylated in murine tissues, with the contribution of chain types to total ubiquitin being ~1% in most tissues (Figure 2a). The lung had ~1.4% of total ubiquitin in the polyubiquitylated form, whereas muscle had the highest with ~8.7%. In most tissues, K48 was the most dominant chain type (Figure 2b), similar to what we observed in BMDMs. However, we observed differences in ubiquitin chain-linkage composition in heart and muscle. Interestingly, K33 was enriched in heart and muscle, compared to the other tissues (Figure 2b). In order to determine the percentage contribution of the K33 linkage type to the total pool of ubiquitin, we normalised this chain type to total ubiquitin in each tissue. K33 contributed only a small proportion to the total pool of ubiquitin, with 5.3 ± 1.9% in muscle and 3.0 ± 1.3% in heart. We verified that these results were not an artefact of our experimental procedure by using a different extraction buffer before Ub-AQUA-PRM analysis. We observed similar results in muscle when using TCA extraction buffer in an independent experiment, with K33 contributing to 5.7 ± 0.9% of the total pool of ubiquitin. Noteworthy, we observed a slightly lower contribution of K33 to the total ubiquitin chain-linkage composition when using TCA extraction, but K33 still contributed >56% to the total ubiquitin chain-linkage composition in muscle (Supplementary Figure S6). This discrepancy is likely due to the different extraction buffer used, which could affect the protein composition extracted from the tissues.

**Figure 2.**
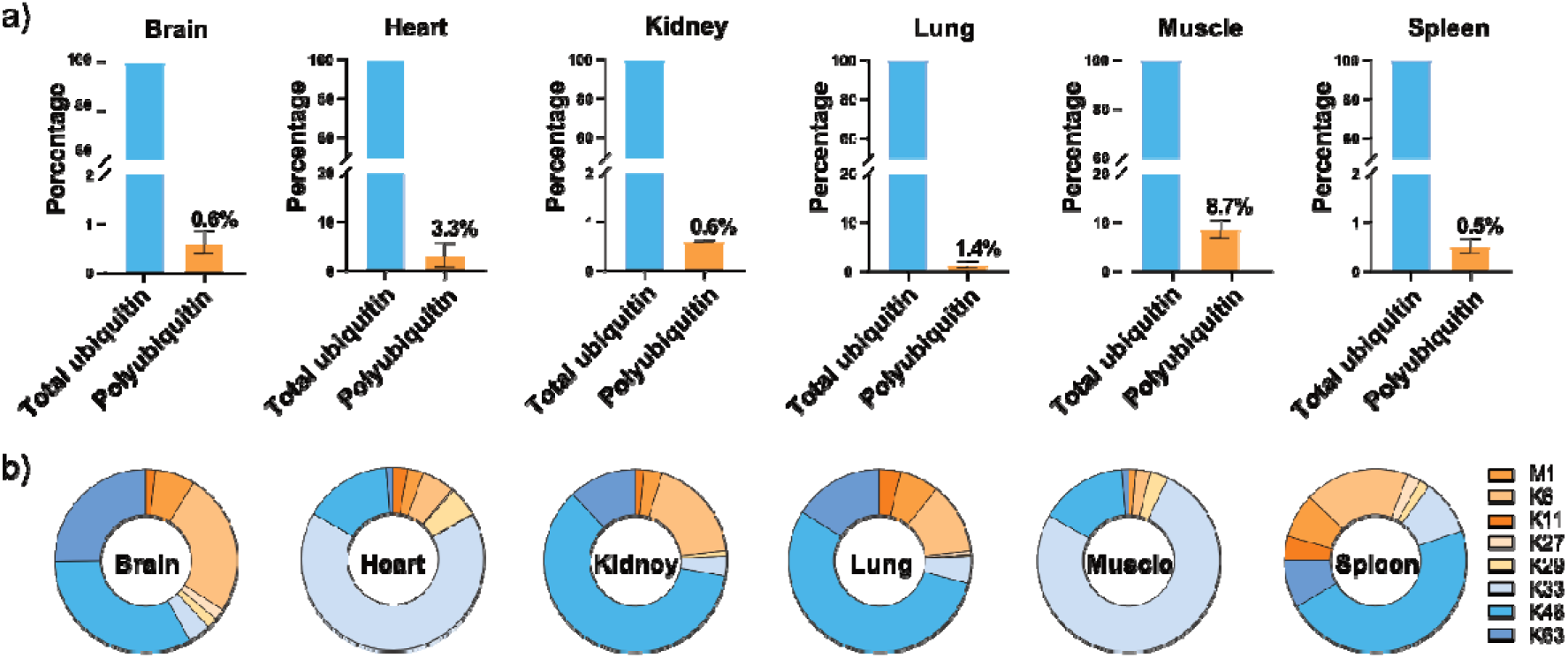
Tissue-specific differences in ubiquitin chain types and compositions exist in murine tissues. (a) Percentage of polyubiquitin chain types present in different murine tissues, compared to the total pool of ubiquitin present in these tissues. (b) Contribution of each of the eight different ubiquitin chain types to the total pool of polyubiquitin in different murine tissues. Error bars indicate standard deviation.

These results indicate that there are tissue-specific differences in total ubiquitin and ubiquitin chain types in murine tissues, with atypical linkages increased in heart and muscle. Furthermore, our data indicate that only a small proportion of total ubiquitin in murine tissues are in the polyubiquitin form. It is, therefore, possible that many proteins in murine tissues are either monoubiquitylated, or that the majority of ubiquitin is in its unconjugated, free form. To test this, we performed immunoblots for total ubiquitin (conjugated and unconjugated) using the same tissues analysed by Ub-AQUA-PRM. We observed high levels of unconjugated ubiquitin in the majority of tissues (Supplementary Figure S7). However, we did observe conjugated ubiquitin in all tissues. More unconjugated ubiquitin was observed in the brain, kidney, lung, and spleen, whereas heart and muscle had the least amount of unconjugated ubiquitin. In our Ub-AQUA-PRM assay, we also observed lower levels of total ubiquitin in the heart and muscle, which corresponds to the data obtained with the western blots. This data indicates that there are major tissue-specific differences in ubiquitylation and that only a small proportion of total ubiquitin in murine tissues are in the conjugated form. Further analysis of the ubiquitylome and ubiquitin chain-linkage landscape, in combination with recently described methods such as Ub-clipping [21] and Ubiquitin Chain Enrichment Middle-Down Mass Spectrometry (UbiChEM-MS) [23], could reveal detailed information on atypical ubiquitylation and higher-order chain-linkage architecture in different tissues and pathological states.

## Methods – online methods

### Preparation of AQUA peptides

Synthetic ubiquitin peptides were obtained from Cambridge Research Biochemicals (Cleveland, United Kingdom) (Supplementary Table S1). Methionine-containing peptides were synthesised under their reduced form, and were subsequently oxidised during sample preparation, as described below. Peptides were dissolved at a concentration of 1 nmol/μl in 50:50 acetonitrile:water (v:v) and peptide purity was monitored by LC-MS/MS. Peptides lacking net peptide content were injected together with their isotope counterpart and net peptide content was determined based on the ratio between the two isotopes. Two sets of master mixes were prepared, with the 10 light and 10 heavy peptides mixed, separately, in equimolar concentrations. The light and heavy peptide master mixes were diluted, separately, to 10 pmol/μl in HPLC-grade water (Thermo Fisher Scientific) and stored at −80°C.

### Optimization of methionine oxidation and ion-pairing agents for targeted proteomic analysis of Ubiquitin AQUA peptides

Methionine oxidation was optimised to generate a stable methionine sulfone for targeted proteomic analysis. Heavy and light peptides were mixed in a 1:1 ratio and incubated with different concentrations of H_2_O_2_ at either room temperature (RT) or 60°C. The conversion of reduced methionine to sulfoxide and sulfone derivatives were monitored by LC-MS/MS. The effect of ion-pairing agents on peptide retention and intensity was assessed by using increasing concentrations of formic acid (FA) and trifluoracetic acid (TFA) in the sample and loading buffers, as indicated in the text.

### Preparation of standard curves

The heavy master mix was diluted to 100 fmol/μl and the light master mix was serially diluted from 10 pmol/μl to 1 amol/μl using a dilution factor of √10. The standard curve was prepared on ice by combining reaction components in the following order: 20 μl of HPLC-grade water, 10 μl of heavy mix at 100 fmol/μl, 10 μl of light mix at varying concentrations, 50 μl of 10% FA, and 10 μl of 10% H_2_O_2_. A second standard curve was prepared by substituting 10 μl of HPLC-grade water with 10 μl of *Escherichia coli* tryptic digest at 1 μg/μl. The *E. coli* tryptic digest was generated as described below. Samples were incubated for 2h at 60°C for oxidation, before targeted proteomic analysis.

### Bacterial cell culture for proteomic analysis

*E. coli* DH5-α was cultured overnight in Luria-Bertani (Thermo Fisher Scientific) broth at 37°C with shaking at 180 rpm. The next day, cells were harvested (4,000 xg, 15 min, 4°C), washed twice with ice-cold PBS pH 7.4 and stored at −80°C until further analysis. Cell pellets were lysed in 5% SDS in 50 mM triethylammonium bicarbonate (TEAB) pH 7.5, by sonication. Whole-cell lysates were subsequently centrifuged (10,000 xg, 5 min, RT) and the supernatants used for proteomic sample preparation.

### Murine cell culture and tissue collection

Animal work was approved by the Newcastle University ethical committee and performed under UK Home Office project licence (PCI 23A338). Bone marrow-derived macrophages (BMDMs) were derived from femurs of 8 to 12-week-old C57BL/6 mice. Bone marrow was extracted and differentiated into macrophages using Iscove’s Modified Dulbecco’s Media (IMDM; Sigma-Aldrich) supplemented with 10% heat-inactivated foetal calf serum (Thermo Fisher Scientific), 1% L-glutamine (Lonza), 1% penicillin/streptomycin (Lonza) and 20% L929-conditioned media for 7 days at 37 °C in 5% CO_2_ humidified atmosphere. After differentiation, BMDMs were stimulated with 10 μM MG132 (Sigma-Aldrich) for 6 hours or mock treated. After stimulation, BMDMs were washed twice with ice-cold PBS pH 7.4. Cells were subsequently lysed by sonication in 5% SDS in 50 mM TEAB pH 7.5, supplemented with 100 mM N-ethylmaleimide (NEM, Sigma-Aldrich) to inhibit deubiquitylases. Whole-cell lysates were used for proteomic sample preparation.

Tissues (brain, heart, lung, kidney, muscle and spleen) were collected from 8 to 12-week-old C57BL/6J mice, washed once with ice-cold PBS pH 7.4, flash frozen in liquid nitrogen and stored at −80°C until further analysis. Tissues were ground to a powder in liquid nitrogen using a mortar and pestle, and lysed by sonication in lysis buffer (5% SDS in 50 mM TEAB pH 7.5) supplemented with 100 mM NEM. Tissue lysates were subsequently centrifuged (10,000 xg, 5 min, RT) and the supernatants used for methanol chloroform precipitation. Protein pellets were resuspended in lysis buffer and used for proteomic sample preparation. In a separate experiment, muscle samples were ground to a powder in liquid nitrogen using a mortar and pestle, and proteins extracted in 20% trichloroacetic acid (TCA) at 4°C. Samples were centrifuged (10,000 xg, 10 min, RT) and washed twice with ice-cold acetone to remove remaining TCA. Protein precipitates were subsequently resuspended in lysis buffer, containing 100 mM NEM, centrifuged to remove debris, and processed as described below.

### Proteomics sample preparation

Protein concentration for all samples was determined using the bicinchoninic acid assay (BCA) assay (Thermo Fisher Scientific). Fifty μg of total protein was used for proteomics sample preparation by suspension trapping (S-Trap) [24,25], with minor modifications to that recommended by the supplier (ProtiFi, Huntington NY, USA). Samples were reduced with 5 mM tris(2-carboxyethyl)phosphine (Pierce) for 15 min at 37°C, and subsequently alkylated with 10 mM NEM for 15 min at room temperature in the dark. Five μl of 12% phosphoric acid was added to each sample after reduction and alkylation, followed by the addition of 330 μl S-Trap binding buffer (90% methanol in 100 mM TEAB pH 7.1). Acidified samples were added, separately, to S-Trap mini-spin columns and centrifuged at 4,000 xg for 1 minute. Each S-Trap mini-spin column was washed with 400 μl S-Trap binding buffer and centrifuged at 4,000 xg for 1 minute. This process was repeated for five washes. Twenty-five μl of 50 mM TEAB pH 8.0 containing sequencing-grade trypsin (1:10 ratio of trypsin:protein) was added to each sample, followed by proteolysis for 2 hours at 47°C using a thermomixer (Eppendorf). Peptides were eluted with 50 mM TEAB pH 8.0 and centrifugation at 1,000 xg for 1 minute. Elution steps were repeated using 0.2% FA and 0.2% FA in 50% acetonitrile, respectively. The three eluates from each sample were combined and dried using a speed-vac before storage at −80°C. Peptides were subsequently reconstituted to 1 μg/μl using 5% FA in water and prepared for PRM, on ice, by combining reagents in the following order: 30 μl of HPLC-grade water, 10 μl peptide digest at 1 μg/μl, 10 μl of heavy master mix at 100 fmol/μl, 40 μl of 10% FA and 10 μl of 10% H_2_O_2_. Samples were incubated for 2h at 60°C for oxidation before targeted mass spectrometry analysis.

### Targeted LC-MS/MS analysis - Parallel Reaction Monitoring (PRM)

Peptide samples were injected on a Dionex Ultimate 3000 RSLC (Thermo Fisher Scientific) connected to a Q-Exactive HF mass spectrometer (Thermo Fisher Scientific). Fifty fmol of each ubiquitin peptide was injected during analysis, along with 500 ng of tryptic peptide mixture from either *E. coli* or *Mus musculus* as indicated in the text. In order to minimise dead volume and shorten the analysis to only 10 min, we used a direct injection LC setup. During the loading phase, samples are loaded using a 5 μl loop and full loop injection, with a 2.4 μl needle and an overfill ratio of 1.5. Samples were injected onto a 150 μm × 150 mm EASY-Spray LC column (Thermo Fisher Scientific) using the nano-pump and 100% buffer A (96.9% HPLC-grade water, 3% DMSO, 0.1% FA) at 1.5 μl/min. After 4 min (6 μl), the column oven valve was switched to bypass the loop and peptides were eluted by increased concentration of buffer B (80% HPLC-grade acetonitrile, 16.9% HPLC-grade water, 3% DMSO, 0.1% FA) using a multistep gradient (Supplementary Table S2). The source was heated to 320°C and 2.5 kV was applied between the column emitter and the MS. The Q-Exactive HF was operated in PRM mode using a resolution of 15,000, AGC target of 2 × 10^5^ and maximum injection time of 25 ms. Peptides were selected for MS/MS data acquisition using pre-determined isolation windows and fragmented using collision energies described in Supplementary Table S1.

### Targeted proteomic data analysis

All Ub-AQUA-PRM RAW mass spectrometry files were analysed using Skyline-daily (version 4.1.1.11871) [26]. Fragment ion intensities were extracted and light to heavy (L/H) ratios of peptide pairs were automatically determined in Skyline. All extracted ion chromatograms were manually inspected. Standard curves were generated from log transformed L/H ratios and peptide concentrations, to determine the lower limit of detection (LLOD) and lower limit of quantification (LLOQ) for each peptide. The R^2^ was determined based on the LLOQ of each peptide. Only curves with a R^2^ value > 0.99 were used for determining the absolute concentration of total ubiquitin and the different ubiquitin chain types. Group comparisons were performed using a Student’s t-test, and multiple group comparisons were performed using an analysis of variance (ANOVA), followed by a Tukey’s HSD test for determining significance for pairwise comparisons, in GraphPad Prism (version 8.0.1). The mass spectrometry proteomics data have been deposited to the ProteomeXchange [27] Consortium via the PRIDE [28] partner repository with the dataset identifier PXD017754. Reviewers can access it through the Username: and the password vJ56s8uk.

### Western blots

Total protein from brain, heart, kidney, lung, muscle and spleen were extracted as described above. Ten-microgram protein from each sample were separated on 12% SDS-PAGE gels before being electrotransferred to Immobilon-P transfer membranes (Merck Millipore). Membranes were probed with monoclonal antibody for mono-and polyubiquitylated proteins (FK2)(BML-PW8810; Enzo Life Sciences), anti-α-Tubulin (T9026; Sigma-Aldrich) and anti-Histone H3 (D1H2) (4499; Cell Signaling) antibodies. Corresponding secondary antibodies used included anti-mouse IgM, HRP-conjugated (AP307P; Sigma-Aldrich) and anti-rabbit IgG, HRP-conjugated (7074; Cell Signaling Technology) antibodies. Peroxidase activity was developed using the Amersham™ ECL™ Prime Western Blotting Detection Reagent (GE Healthcare) and imaged on an Amersham Imager 600 digital imaging system (GE Healthcare). To assess equal loading of protein, Ponceau staining (Sigma-Aldrich) of the PVDF membranes was performed.

## Acknowledgements

This work was funded by a Wellcome Trust Investigator Award (215542/Z/19/Z) and the generous start-up funding of Newcastle University to MT. FL was funded by a Marie Skłodowska-Curie Fellowship from the Horizon 2020 programme of the EU (project number 795670).

## Conflict of interest

The authors declare no conflict of interest.

## Author contributions

TH, FL and MT designed the study. TH and FL performed mass spectrometry experiments and data analysis. JLM-R performed western blots. AD was responsible for animal husbandry and animal experimental procedures. TH and MT wrote the paper with contribution of all authors.

## Supporting documents

**Supplementary Figure S1.**
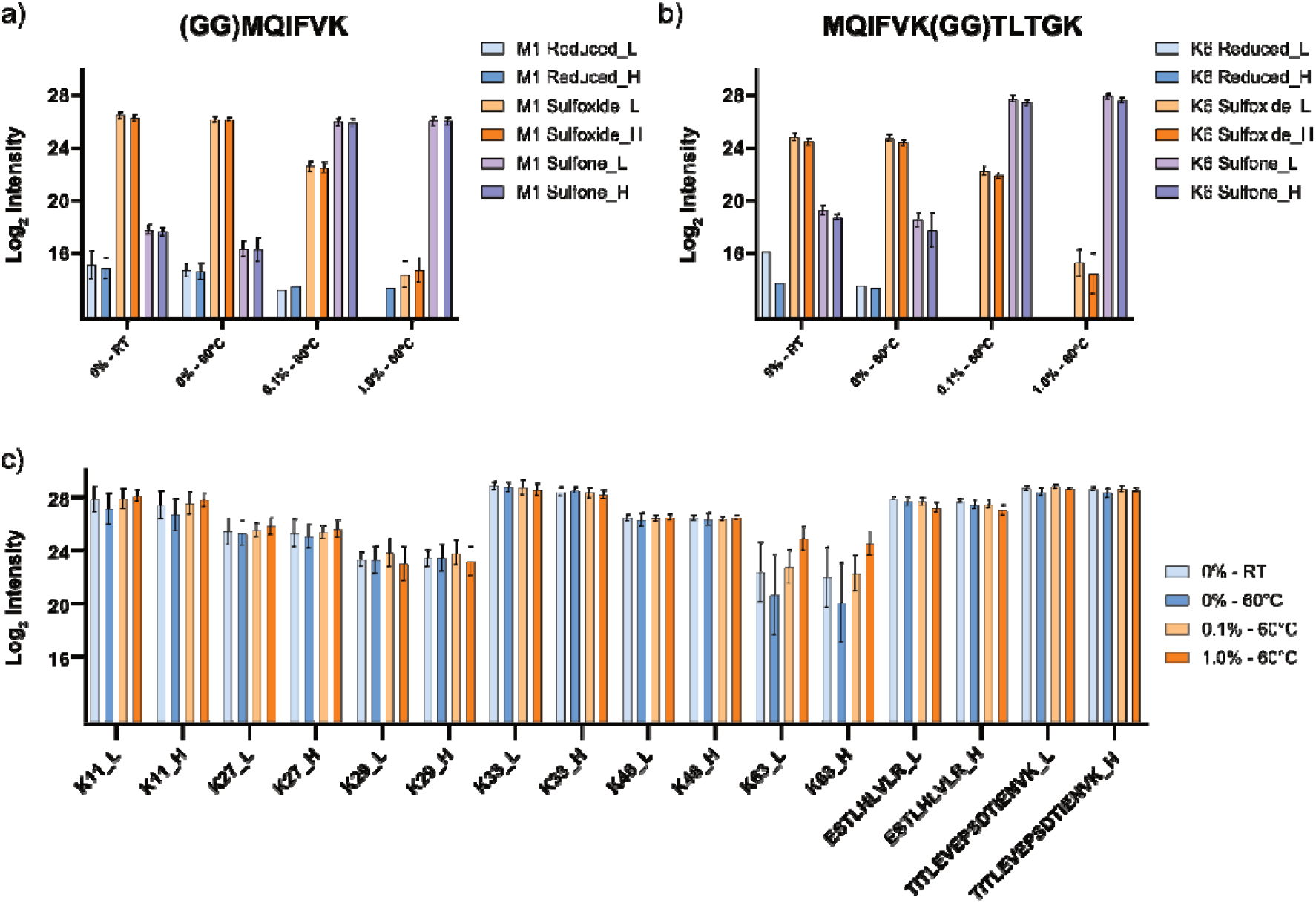
Generation of a stable methionine sulfone by oxidation. Methionine-containing ubiquitin peptides were oxidised using hydrogen peroxide (H_2_O_2_) and analysed by LC-MS/MS to determine the generation of a methionine sulfone. A stable methionine sulfone was generated using 1.0% H_2_O_2_ at 60°C for 2 hours in M1 (a) and K6 (b) peptides. (c) Peptide intensities observed for non-methionine-containing ubiquitin peptides when using H_2_O_2_ for oxidation. L = light version of the peptide, H = heavy version of the peptide, LoD = limit of detection. Error bars indicate standard deviation.

**Supplementary Figure S2.**
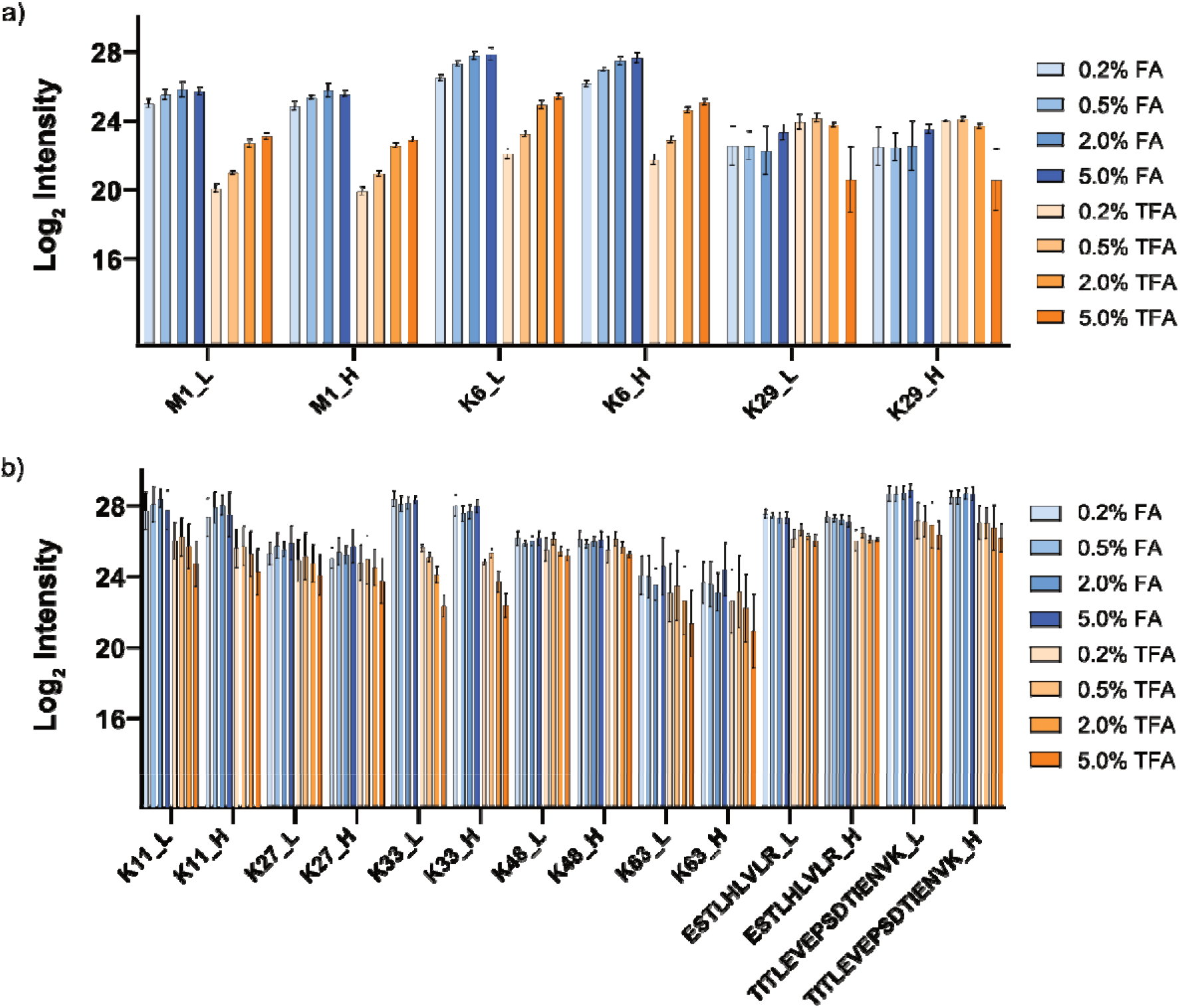
Formic acid is a superior ion-pairing agent for ubiquitin peptides. (a) Peptide intensities observed for M1, K6 and K29 ubiquitin peptides when using formic acid (FA) and trifluoroacetic acid (TFA) as ion-pairing agent in the loading buffer. (b) Peptide intensities observed for the other ubiquitin peptides when using FA and TFA in the loading buffer. L = light version of the peptide, H = heavy version of the peptide, LoD = limit of detection. Error bars indicate standard deviation.

**Supplementary Figure S3.**
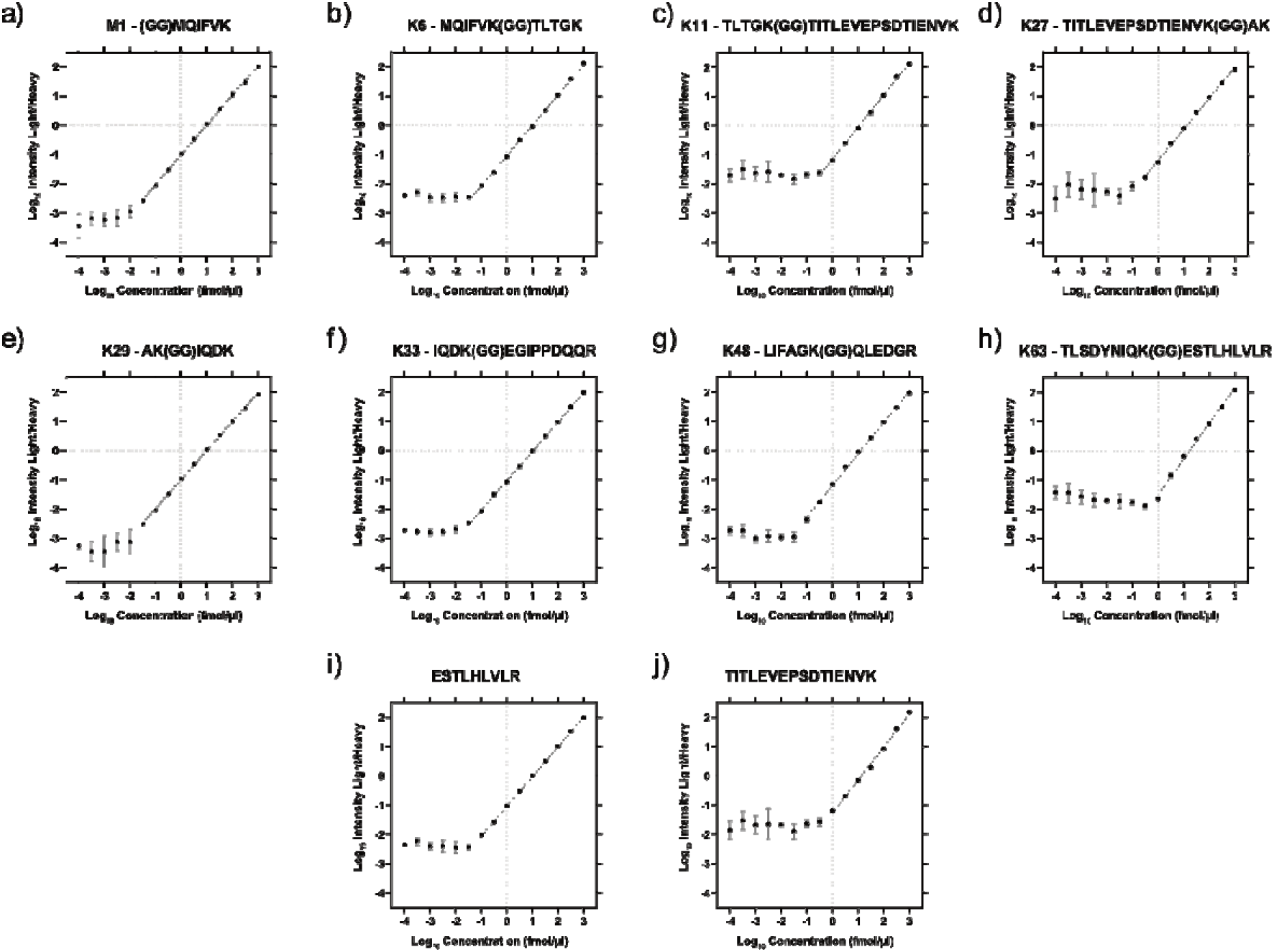
Limits of detection and quantification of spiked-in, heavy and light ubiquitin peptides in an *E. coli* proteome digest. Standard curves were generated using 10 synthetic ubiquitin peptides in their light and heavy counterparts. Heavy peptides were used at 10 fmol/μl, while light peptides were serially diluted from 1 pmol/μl to 1 amol/μl. Peptides were oxidised using H_2_O_2_ before injected on a Q-Exactive HF mass spectrometer for PRM analysis. The peptide master mix contained each of the eight different ubiquitin chain-type peptides (a-h) and two peptides that served as proxy for total ubiquitin (i, j). Error bars indicate standard deviation.

**Supplementary Figure S4.**
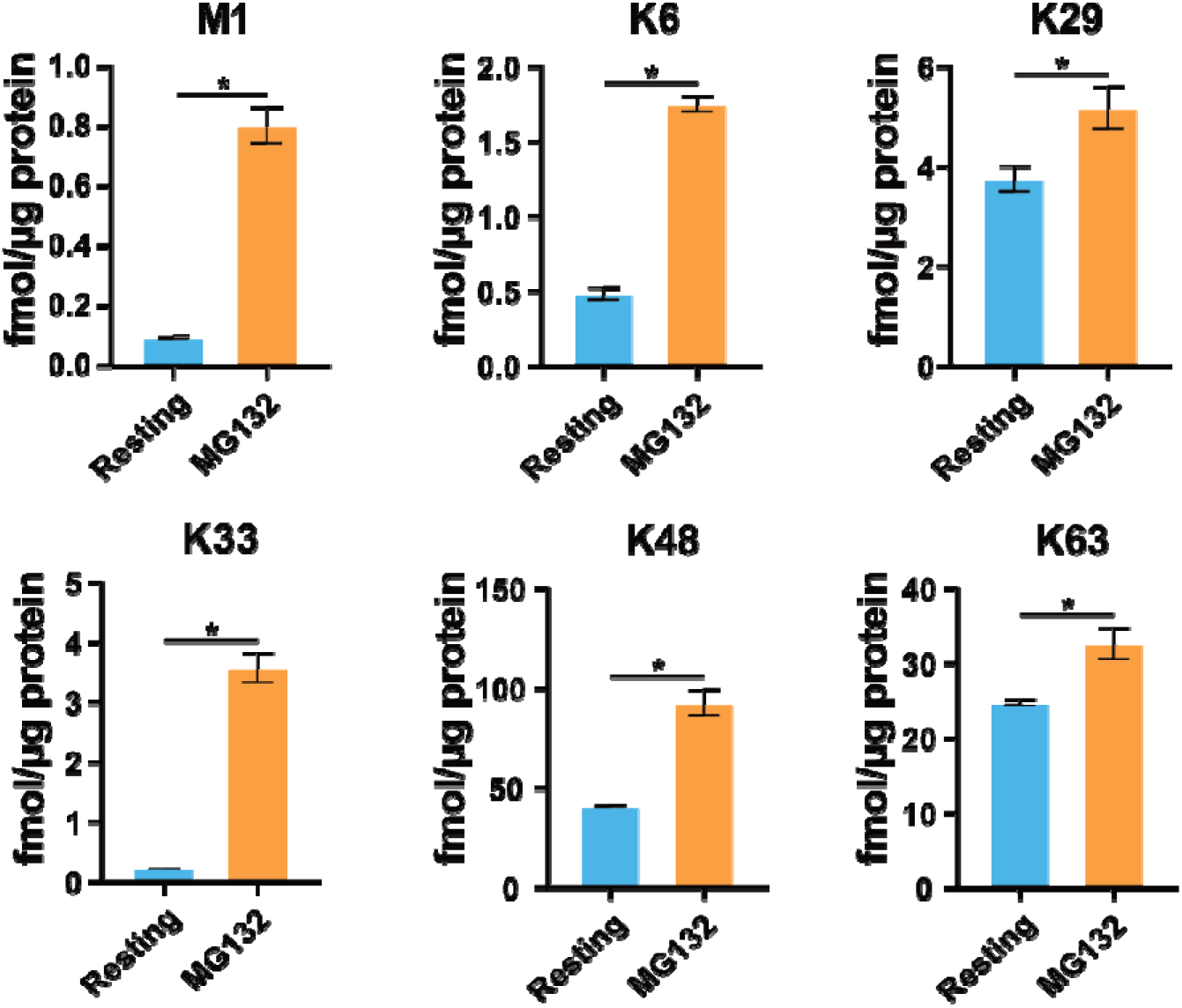
MG132 treatment causes an increase in ubiquitin chain types. Differences observed in the quantified ubiquitin chain types at resting state and during MG132 treatment. Statistical significance is denoted with an * and indicates a p-value < 0.05 determined using a Student’s t-test. Error bars indicate standard deviation.

**Supplementary Figure S5.**
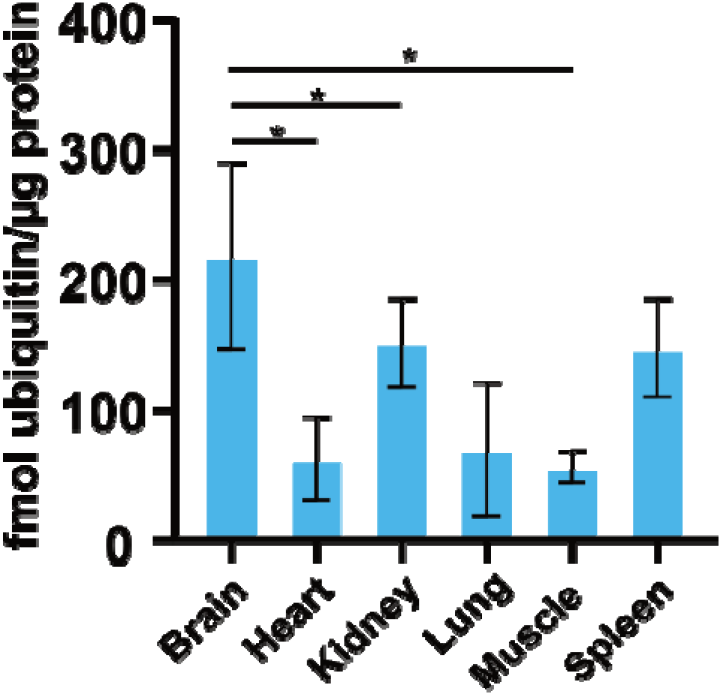
Murine tissues contain different levels of ubiquitin. Absolute concentration of ubiquitin in different murine tissues when using non-modified peptides as proxy for total ubiquitin. Total ubiquitin was derived from the sum of the absolute concentrations of TITLEVPSDTIENVK and K11 ubiquitin peptides. Statistical significance is denoted with an * and indicates an adjusted p-value < 0.05 after a Tukey’s HSD test and ANOVA. Error bars indicate standard deviation.

**Supplementary Figure S6.**
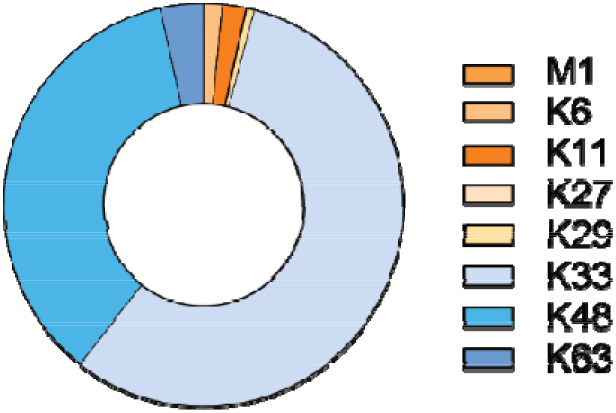
Mouse muscle is enriched in K33 ubiquitin chain types. (a) Contribution of each of the eight different ubiquitin chain types to the total pool of polyubiquitin in mouse muscle when using 20% TCA for protein extraction to inhibit deubiquitylase activity.

**Supplementary Figure S7.**
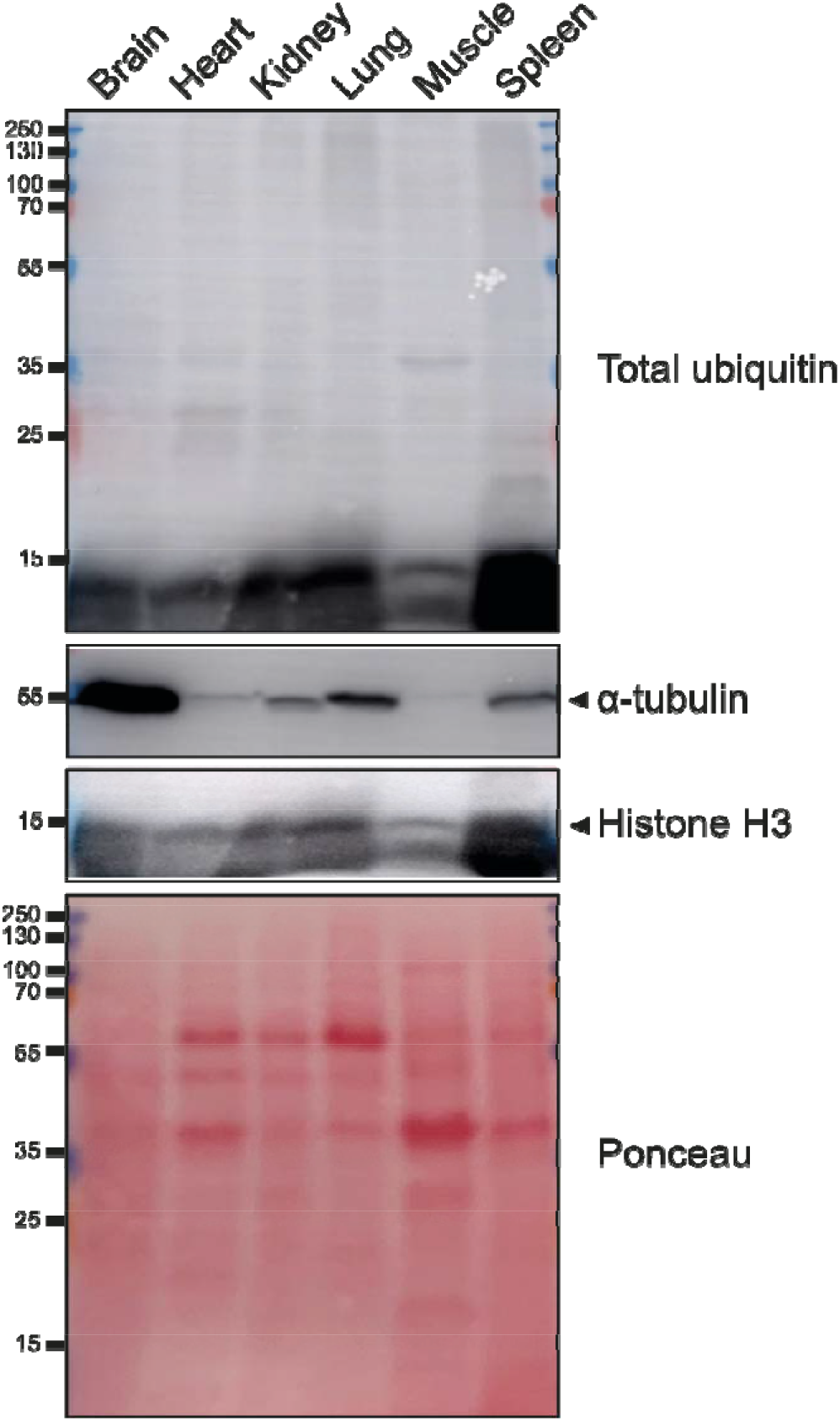
Murine tissues contain a large amount of unconjugated ubiquitin. Western blot for total ubiquitin (conjugated and unconjugated) in murine tissues. The same tissues were used as those analysed by Ub-AQUA-PRM.

**Supplementary Table S1.**
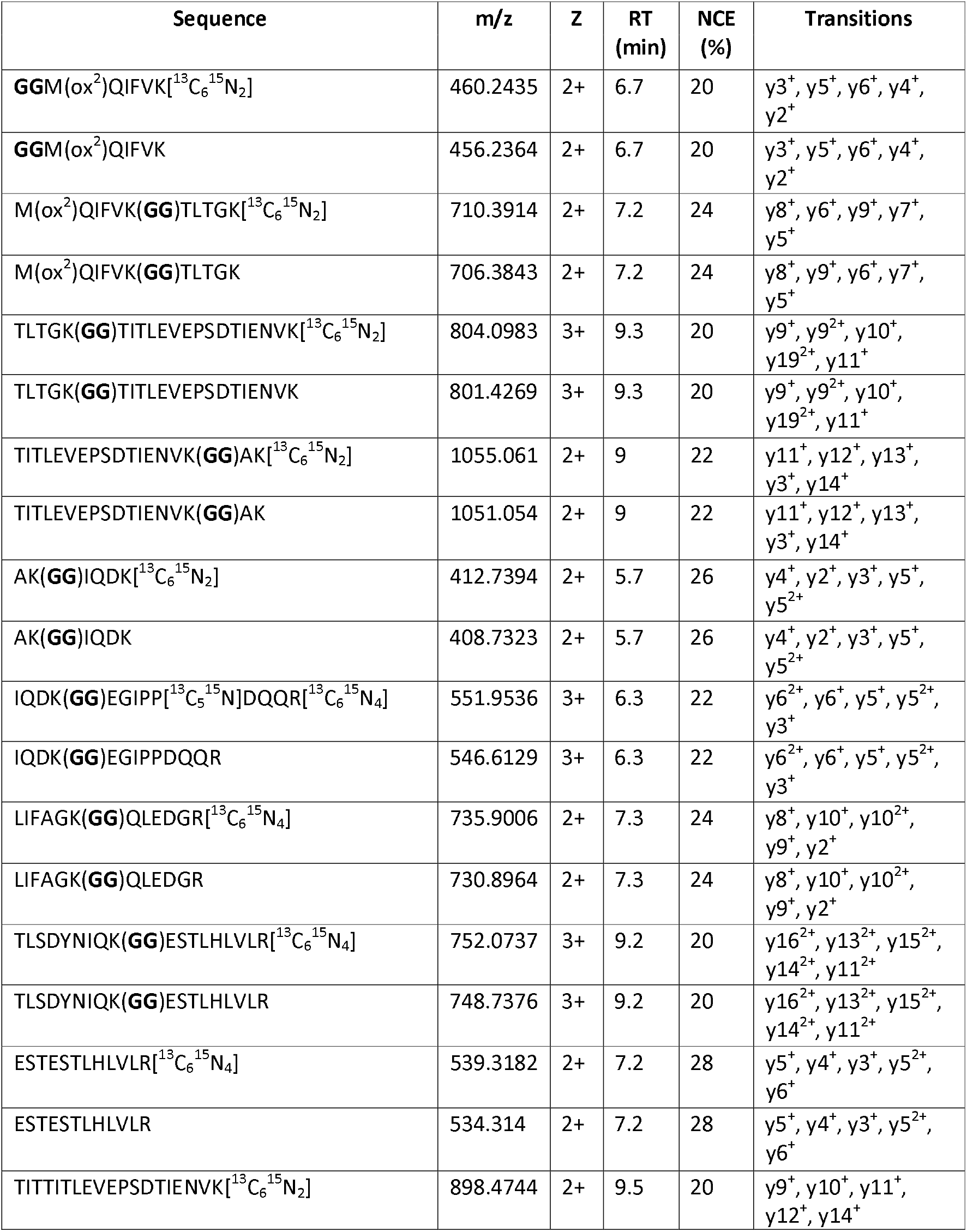
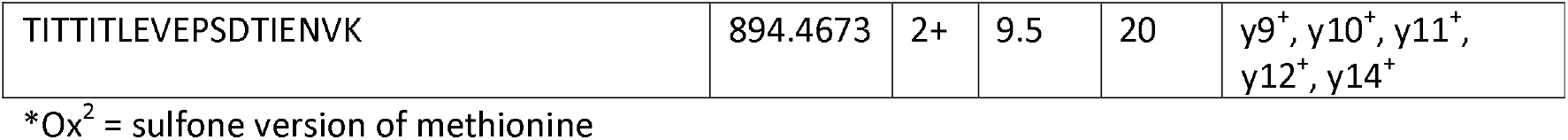
Peptides used for absolute quantification of ubiquitin chain types by Ub-AQUA-PRM analysis.

**Supplementary Table S2.** Multi-step gradient used for separating ubiquitin peptides during Ub-AQUA-PRM analysis.

| Time (min) | %B |
| --- | --- |
| 0 | 0 |
| 4 | 0 |
| 4.25 | 7 |
| 5 | 12 |
| 6 | 15 |
| 7 | 16 |
| 8 | 24 |
| 8.25 | 35 |
| 8.5 | 90 |
| 8.75 | 90 |
| 9 | 0 |
| 10 | 0 |

## Notes

### Competing Interest Statement

The authors have declared no competing interest.

### Summary of Updates

the manuscript has been shortened.

